# Individual variability of neural computations in the primate retina

**DOI:** 10.1101/2021.02.14.431169

**Authors:** Nishal Shah, Nora Brackbill, Ryan Samarakoon, Colleen Rhoades, Alexandra Kling, Alexander Sher, Alan Litke, Yoram Singer, Jonathon Shlens, E.J. Chichilnisky

## Abstract

Variation in the neural code contributes to making each individual unique. We probed neural code variation using ∼100 neural population recordings from major ganglion cell types in the macaque retina, combined with an interpretable computational representation of individual variability using machine learning. This representation captured individual variation and covariation in properties such as nonlinearity, temporal dynamics, and spatial receptive field size, while preserving invariances, such as asymmetries between ON and OFF cells. The covariation of response properties in different cell types was associated with the proximity of lamination of their synaptic inputs. Surprisingly, male retinas exhibited higher firing rates and faster temporal integration than female retinas. Exploiting data from previously recorded macaque retinas enabled efficient characterization of a new macaque retina, and of a human retina. Simulations indicated that combining a vast dataset of healthy macaque recordings with behavioral feedback could be used to identify the neural code and thus improve retinal implants for vision restoration.

## Introduction

An emerging frontier in biomedicine is understanding variability between individuals, with implications ranging from the mathematical modeling of living systems to ethics and personalized medicine. In neuroscience, differences in mental function between individuals are substantial, yet little is known about the underlying variation in the information processing performed by neural circuits, particularly in species similar to humans and at the spatial and temporal scales of neural computations. Furthermore, biological variability is frequently obscured by inevitable experimental variability, which can severely limit the ability to decisively test biological hypotheses. These factors have led to a large gap in our understanding of variability in the neural code and its implications for translational medicine and neuroengineering.

Two technical challenges have limited our understanding in animal models relevant to humans: high-resolution, large-scale physiological recordings from many individuals, and methods for deciphering variability in complex circuit level computations. In this paper, we present a novel method for analyzing variability in the neural code -- both biological and experimental -- and apply it to a unique large-scale dataset gathered over a decade of recordings in the macaque retina. The results reveal striking differences in male and female retinas, structure in variability related to lamination of dendrites, and novel implications for neuroengineering to replace retinas damaged by disease.

## Results

### Modeling the shared features and individual variability of neural coding

Large-scale multi-electrode recordings from retinal ganglion cells (RGCs) were performed from isolated macaque monkey retinas, in which the functional properties of diverse cell types have been extensively studied (Field et al., 2007; Frechette et al., 2005; Greschner et al., 2014; Litke et al., 2004; Rhoades et al., 2019). In total, 21,626 RGCs were analyzed in 112 recordings from 75 retinas of 66 animals. These data, gathered over a decade of experimentation, exhibited significant neural coding variability across recordings, presumably reflecting a mixture of biological and experimental variation (see below). As a baseline to summarize this variability, response properties in each recording were captured by the parameters of a linear-nonlinear-Poisson (LNP) encoding model. This widely used model (Chichilnisky, 2001) captures light-evoked responses in RGCs using a spatiotemporal linear filter applied to the stimulus, followed by an output nonlinearity and stochastic spike generation. Receptive field sizes of two known RGC types -- ON and OFF parasol -- exhibited substantial variation across recordings. This variability was evident across retinal eccentricities, but was also present at a given eccentricity (Figure 1A, first column, Figure 1G). Diversity was also seen in the kinetics of light responses (Figure 1A, second column), the shape of the output nonlinearity (Figure 1A, third column) and the autocorrelation function (Figure 1A, fourth column).

**Figure 1.**
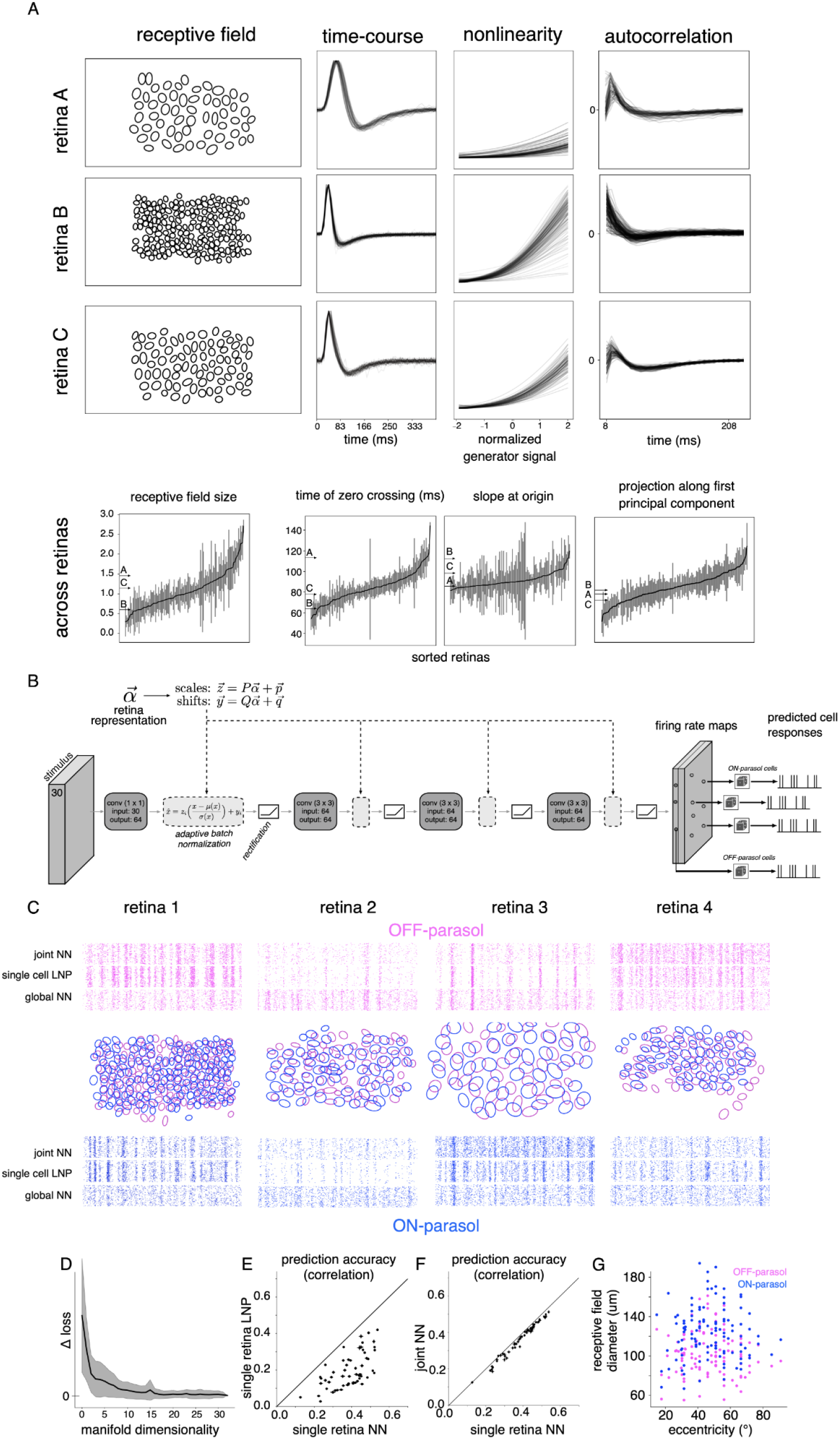
Modeling variability in the neural code. (A) Variability of response properties across recordings. Spatial receptive fields (first column), time of zero-crossing of temporal filter (second column), nonlinearity (third column) and variation in autocorrelation function (fourth column) for OFF parasol cells in three representative recordings (rows, same y-axis for each response property across recordings). The last row represents the range of response parameters across 122 recordings, sorted according to their population means, and error bars corresponding to the robust standard deviation. Arrows indicate the values of the chosen retinas in the first three rows. (B) Architecture of the neural network for capturing response variation. The visual stimulus is passed through multiple layers of convolution with spatial filters, with adaptive batch normalization and rectification at each layer, producing two firing rate maps (one each for ON and OFF parasol cell types). The Poisson firing rate for each cell is read off from the value at the cell’s location in the firing rate map of its cell type. Retina-specific tuning of responses is performed by adjusting the mean and standard deviation of the activation values at each layer, determined by a linear transformation of the retina’s location in the low-dimensional manifold. (C) Response prediction across 4 representative training retinas (columns). The receptive field mosaics are shown for each retina (middle row), along with response predictions for a randomly selected OFF parasol (top row) and ON parasol cell (bottom row). Rasters (60 trials) for predicted responses to a 3 sec long white noise stimulus using the LNP model; neural network model, trained jointly on multiple retinas, with retina-specific parameters (15 dimensional manifold, joint NN) or without them (no manifold, global NN). (D) Model error (log likelihood) on test stimuli with varying manifold dimensionality; dimensions=0 indicates no retina-specific adaptation (global NN). Error bars indicate the standard deviation across retinas. (E) Prediction accuracy, averaged across cells, for different retinas (points), using a neural network trained on data from each individual retina (x-axis) and an LNP model with shared parameters across cells of a given type in each retina (y-axis). Prediction accuracy is measured as correlation between predicted firing rate and recorded responses smoothed with a Gaussian filter (σ: 11ms). (F) Similar to (E), comparing predictions from the neural network with a 15-dimensional manifold, trained jointly (y-axis) vs. trained on each retina separately (x-axis). (G) Range of eccentricities (x-axis) and the average receptive field sizes (y-axis) for ON parasol (blue) and OFF parasol (magenta) cells across 122 recordings used in this study.

The structure and covariation of these response properties was explored using a flexible model that combined shared and recording-specific parameters. The shared component was a multilayered convolutional neural network (CNN), an extension of the LNP model, consisting of multiple alternating stages of spatio-temporal filtering, normalization and rectification. The rectification captured nonlinear spatial integration (Shah et al., 2020), and the convolutional structure captured the known translational invariance of visual signals in each cell type (cells of the same type at different locations have very similar response properties (Chichilnisky and Kalmar, 2002). The model output consisted of one firing rate map for each cell type. To predict a given cell’s responses, the model-predicted firing rate was read off from the map at the cell’s location (Figure 1B). Due to the translational invariance constraint, the proposed model cannot capture differences between cells belonging to the same cell type, resulting in low prediction accuracy compared to models that allow for cell-specific parameters, such as single-cell LNP (Chichilnisky, 2001) or other state-of-the-art models (Batty et al., 2017; McIntosh et al., 2016). However, the translational invariance constraint enabled the proposed neural network architecture to predict responses across recordings with different numbers and spatial arrangements of cells. When trained using ON and OFF parasol cell responses in each retina separately, the CNN model exhibited performance superior to the single-retina LNP model with common parameters for all cells of a given type, as expected given its more flexible structure (Figure 1E). However, when the CNN model was trained on multiple retinas together, it failed to capture the responses of retinas with low firing rates, highly modulated responses, or other features that varied between recordings (Figure 1C, rasters).

To capture the variation of light response properties across recordings in a compact and tractable way, data from all recordings were used to learn the ∼100K parameters of the shared CNN, while data from each recording were used to estimate a small number of recording-specific parameters (see below). A linear transformation of the recording-specific parameters determined the mean and standard deviation of signals in the channels in each layer of the CNN. The collection of these recording-specific parameters was interpreted as a *manifold of neural coding variability*. When learned using 71 recordings, the low-dimensional manifold captured variations in background firing rate, sustained vs. transient dynamics, and response nonlinearities (Figure 1C, rasters). The ability to simultaneously predict responses across multiple retinas saturated at ∼15 dimensions of the learned manifold (Figure 1D), much lower than the total number of CNN parameters, and the performance of the joint model based on the manifold was only slightly lower than that of a CNN model trained for each retina separately (Figure 1F). Thus, a simple, low-dimensional representation can efficiently and accurately capture the biological and experimental variability of retinal computation.

### Neural coding manifold smoothly captures systematic variation across recordings

The learned manifold smoothly captured variations in the neural code. For a given recording, a greater perturbation in manifold location led to a greater decrease in response prediction accuracy (Figure 2G). The manifold geometrically represented variation in several light response properties, including firing rate, receptive field size, time course, output nonlinearity, and spike train autocorrelation. This was observed by projecting the average response property for each retina onto its principal component across all recordings, and then identifying the manifold direction with maximum correlation to this projection. The Spearman rank correlation between the principal component projections and the projections along the identified manifold direction was significantly higher than the value observed in random permutations of the data (p < 0.05 for all response properties) (Figure 2B-F).

**Figure 2.**
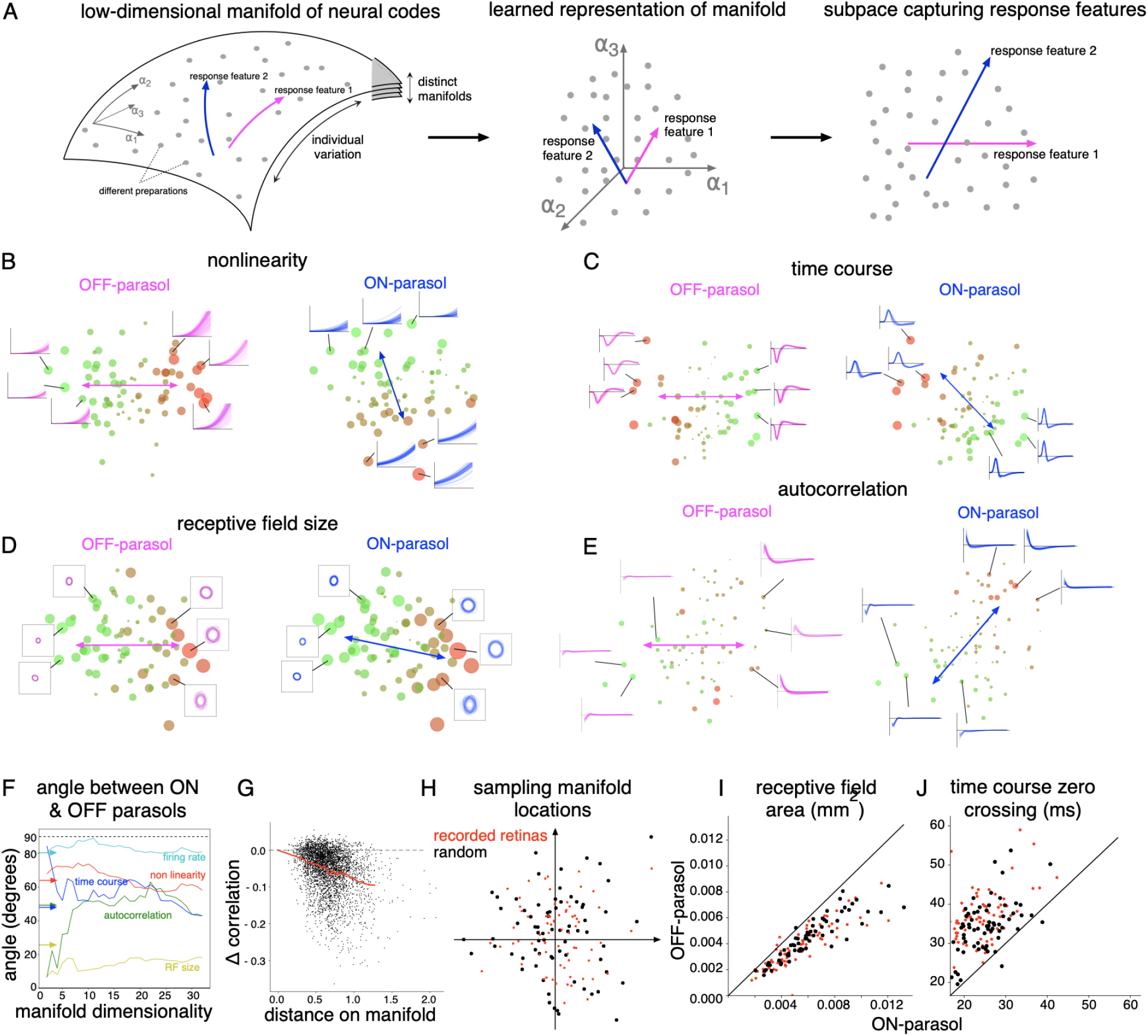
Geometry of manifold. (A) Summary of the steps in subsequent analysis. Left: Schematic of the manifold representation of variability. Each recording is summarized by its neural encoding function, indicated by a point (gray) in space of all possible neural encoding functions. The observed neural encoding functions lie in a low-dimensional manifold (depicted as curved surface, but could be more than two dimensional) within this space. Different manifolds (other surfaces) would potentially correspond to different properties that are invariant across recordings. The training procedure learns a coordinate system (*α*)within the manifold. Middle: Directions corresponding to response features can be identified in the learned coordinate system for representing the manifold. Right: Geometry of the subspace corresponding to the identified directions lead to interpretation of the variations. (B) Manifold locations for 95 recordings (points), projected onto the 2D subspace given by the first principal components of variation in the output nonlinearity for OFF and ON parasol cells. Size and color of dots indicate deviation from the mean. Colored lines indicate the direction of maximum nonlinearity variation for ON parasol (blue) and OFF parasol (magenta) cells. Insets: Lines show output nonlinearities for all cells in representative retinas. (C, D, E) Similar to (B), for time course, receptive field size and autocorrelation, respectively. (F) Angle between ON and OFF parasol directions for particular response properties, as a function of manifold dimension. Arrows indicate the cosine inverse of Spearman rank correlation computed directly between the response properties. (G) Change in response prediction accuracy (y-axis) as the manifold location is perturbed from the learned location. Each black dot represents a different perturbation, the red line is the average. (H) Random manifold locations (red) were sampled by adding noise to the learned retina-specific locations (black). Responses to a white noise stimulus of 100 cells of each type (ON and OFF parasol) were sampled from random locations in the visual field using the firing rate maps for the two cell types associated with these sampled manifold locations. These simulated responses were then used to fit a LNP model. (I, J) Relationship between ON and OFF parasol cells for receptive field area and zero crossing of response time course, respectively, for recorded (red) and randomly sampled (black) retinas

The geometry of the manifold also captured co-variation in response properties of different cell types across recordings. For both firing rate and response nonlinearity, the large angles between the manifold directions for ON and OFF parasol cells (84° and 66° respectively) were consistent with low Spearman rank correlation in these response properties (0.17 and 0.44 respectively). Conversely, for response autocorrelation, time course and receptive field size, the small angles between the directions for ON and OFF cells (49°, 54° and 13° respectively) were consistent with the larger Spearman rank correlation (0.65, 0.67 and 0.90 respectively). Although high correlation is expected for receptive field size based on variation in the eccentricity of different recordings, the varying degrees of covariance for time-course, autocorrelation, output-nonlinearity and firing rate have not been previously reported. Hence, the manifold presents an intuitive, geometric representation of variation and covariation of response properties.

The manifold also captured known *invariances* in retinal encoding, i.e. properties that were consistent across retinas. To assess whether these invariances were present at many intermediate manifold locations not directly sampled in the experiments, the manifold locations of recorded retinas were perturbed using Gaussian noise with standard deviation equal to the median nearest neighbor separation (Figure 2H). Light responses were then generated using these randomly sampled manifold locations, and the encoding properties were summarized by fitting a LNP model. For both the recorded and simulated responses, OFF parasol cells had consistently smaller RFs (Figure 2I) and slower time courses (Figure 2J) than ON parasol cells, consistent with previously reported asymmetries (Chichilnisky and Kalmar, 2002).

To the degree that these systematic variations represent differences between animals, a simple prediction is that recordings from the same animal should be closer on the manifold than recordings from different animals. The results confirmed this prediction (paired t-test, p<0.01 across manifold dimensions). Note, however, that animal-specific variations in experimental procedures could also produce this result (see Discussion), motivating a deeper look at variations in the manifold that likely reflect true biological differences.

### Manifold reveals covariation associated with retinal connectivity, and male-female differences

The RGC types that receive synaptic input from bipolar cells at similar depths in the inner plexiform layer (IPL) showed greater covariation in their response properties across recordings than other RGC types. To examine covariation between cell types, the ON and OFF midget cell types were included with the ON and OFF parasol cell types considered thus far. In 85 recordings (53 macaques), the similarity of three response properties – firing rate, nonlinearity, and time course – across different pairs of cell types was measured either directly, or in the manifold (Figure 3A-C). Using both methods, the highest correlation in these physiological properties was observed between cell-type pairs with the same polarity (ON or OFF), consistent with the lamination of ON and OFF cells in the inner and outer IPL, respectively (Figure 3D). Moreover, for cell type pairs with opposite polarities, a higher correlation in physiological properties was observed for the ON-parasol/OFF-parasol pair than for the ON-midget/OFF-midget pair, consistent with the lamination of parasol cells closer to the middle of the IPL (Figure 3A-D). These observations support the approach of studying the response properties of newer cell types (such as ON and OFF smooth monostratified cells (Rhoades et al., 2019)) after normalizing to the properties of more commonly studied cells with similar synaptic inputs, thereby minimizing the effects of inter-retina variability.

**Figure 3.**
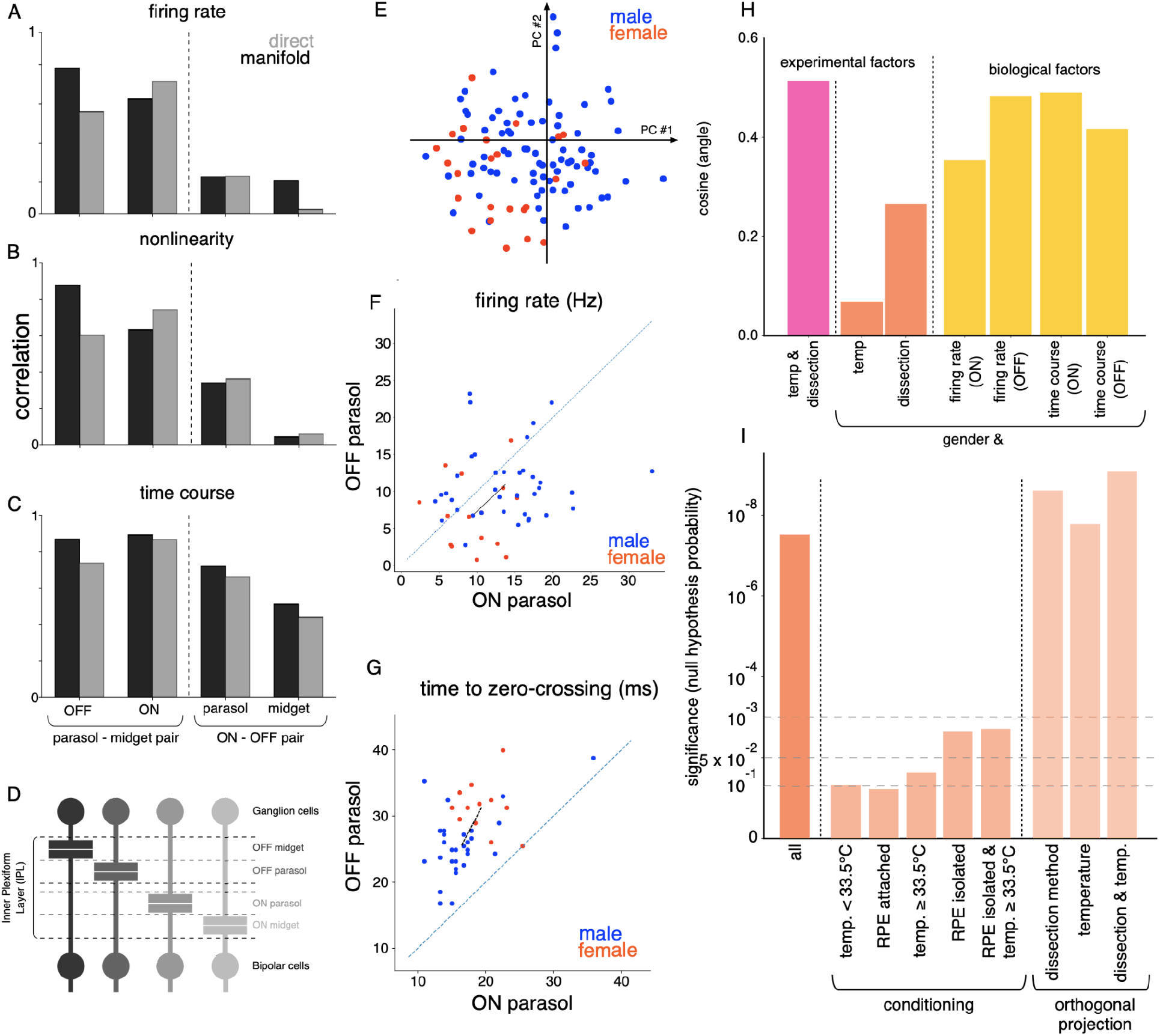
Biological factors underlying variability. Relation between the first principal component of (A) firing rate, (B) nonlinearity and (C) response time course variation for different pairs of cell types. The relationship is either measured directly using Spearman rank correlation, or the cosine of the angle between corresponding directions in the 15-dimensional manifold. (D) Distinct lamination depths for the bipolar-ganglion cell synapse for different ganglion cell types(Wassle and Boycott, 1991). (E) Two dimensional PCA projection of manifold locations for recordings from male (blue) and female (red) retinas. (F) The average firing rate for ON parasol (y-axis) and OFF parasol (x-axis) cells for recordings with isolated RPE. The mean manifold location of male (blue) and female (red) recordings were different (p<0.01 for bootstrap and p<0.05 for hierarchical bootstrap). Black line joins the mean male and female locations. (G) Similar to (F) for the time course of STA, with separation of male and female retinas (p < 0.01 for bootstrap and p<0.05 for hierarchical bootstrap (Saravanan et al., 2019)). (H) Cosine of the angle (y-axis) between the manifold directions corresponding to different pairs of factors, which are either biological (sex, firing rate, time of zero crossing of time course) and experimental (temperature, dissection -whether retinal pigment epithelium (RPE) was attached or isolated). (I) Degree of separation of male and female recordings measured using a resampling test, for all the recordings, conditioned on the subset of recordings with specific dissection procedure or temperature, or all recordings with locations projected orthogonal to directions for dissection procedure and temperature variation.

Surprisingly, recordings from male and female retinas were separated in the manifold (d’=1.8 for 15 dimensional manifold, Figure 3E), in a way that was not explained by variations in experimental factors (Figure 3I). For both ON and OFF parasol cells, the differences between male and female retinas were partially attributable to differences in firing rate and speed of temporal filtering. This was determined by computing the direction separating the means of male and female retinas in the manifold, and determining the angle between this direction and the directions most aligned with variation in firing rate and response time course (cosine(angle) ∼ 0.5 for both) (Figure 3H). In principle, the observed male-female differences could be confounded by variation in experimental methods such as dissection procedure (isolation from the retinal pigment epithelium, or RPE) or temperature (31°-36°C across recordings). Because higher recording temperatures were associated with isolation from the RPE for technical reasons, the directions in the manifold representing dissection method and temperature variation were aligned (cosine(angle) ∼ 0.57, Figure 3H). Compared to firing rate and response time course, these experimental factors were less aligned to the direction of sex separation (cosine(angle) ∼ 0.26) (Figure 3H), suggesting that sex differences were probably not due to differences in these experimental methods. To eliminate experimental factors more rigorously, the separation of males and females in the manifold was measured after conditioning the data in several ways. For each condition, a bootstrap rank test was performed to test if the mean locations of male and female recordings differed (see Methods). Significant separation (p<0.05) was observed for the more numerous recordings with RPE-isolated dissections (37 male, 15 female) and high temperature (≥33.5°C) (32 male, 12 female), whereas the separation was not significant (p>0.1) for less numerous dissections with the RPE attached (12 male, 4 female) and lower temperatures (<33.5°C) (19 male, 5 female) (Figure 3I). For the RPE-isolated retinas, the male recordings exhibited higher firing rates (Figure 3F) and faster temporal integration (Figure 3G) (p<0.01, see Methods for details).

Although the conditioning on specific experimental conditions above revealed statistically significant differences between males and females, the level of significance was lower when compared to all the recordings (Figure 3I), potentially due to a reduction in the number of samples when analysis was restricted to a particular set of experimental conditions. The manifold made it possible to separate experimental variations more efficiently, without reducing the number of data points. To accomplish this, the data were projected onto axes in the manifold orthogonal to the two identified directions of experimental variability, namely dissection method and temperature variation (a separate analysis showed that two directions sufficed to capture these two types of variability; not shown). This projection increased the statistical significance of separation between the male and female retinas (p<10^−6^) (Figure 3I). Thus, the geometry of the manifold makes it possible to examine statistical trends in the data efficiently in spite of potential experimental confounds.

### Manifold generalizes to a novel recording

The manifold permitted efficient response modeling of a new, previously unseen retina by leveraging trends in the large data set of retinas used for learning the model. Response modeling can be performed efficiently by identifying the manifold location of a new retina in several ways, using limited data. In the absence of any new data, the “expected” manifold location for a new retina can be obtained by merely *averaging* the locations of all training retinas. In the case of a degenerated retina (in which no light-evoked response can be recorded), partial information such as the firing rate of recorded neurons can be identified from spontaneous activity. In this case, the manifold location can be *approximated* by averaging the locations of training retinas that have similar firing rate. Finally, in the presence of measured light-evoked responses, gradient descent can be used to *optimize* the manifold location based on the likelihood of the data, leveraging the training retinas as prior information (Figure 4A).

**Figure 4.**
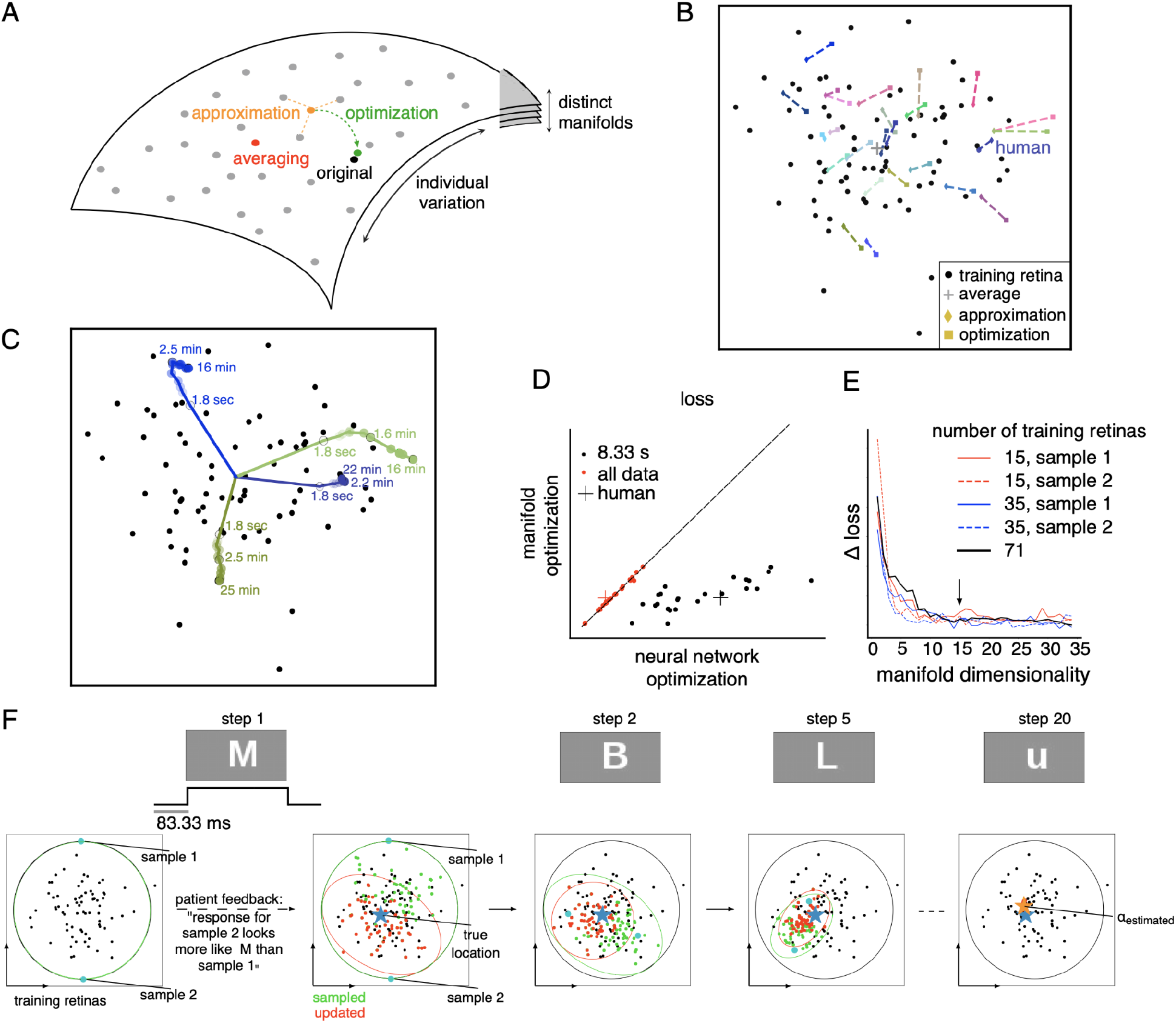
Generalization to new, previously unseen retinas. (A) Efficient identification of manifold location. Training retinas (gray points) are used to define the manifold. Then, the manifold location of a new retina is determined in one of several ways - averaging (red): compute the mean location of all training retinas; approximation (orange): compute the mean location of a subset of training retinas with specific features similar to the new retina; optimization (green): gradient descent on the manifold location to maximize the prediction accuracy for measured light responses of the new retina. (B) Identified manifold locations using averaging (+; same for all retinas), approximation (⋄) and optimization () for testing retinas (colors, each pair joined with a line). Black points correspond to the locations of training retinas. (C) Optimized manifold location for three retinas (colored lines), with varying duration of recorded responses. Training retina locations (black points) and locations identified by averaging (+) are shown. For (B, C), the 15 dimensional manifold is projected into two dimensions that capture firing rate variation (same as Figure 2E) (D) Response prediction loss with optimization of manifold location (y-axis) vs. loss with learning all neural network parameters (x-axis), using either 8.33ms (red) or 15-30 min (black) of data. (E) Convergence of prediction loss with optimized manifold location, averaged across 24 testing retinas (y-axis) as a function of the number of manifold dimensions (x-axis). Loss measured as negative log-likelihood, averaged across cells. Colored lines indicate loss obtained using fewer training retinas. (F) Simulation of the discrimination task used to identify manifold location in the retina of a blind person. At each step, the visual stimulus is a letter from English alphabet (top row). Two dimensional projection of the manifold with locations of training retinas (black dots) is shown (bottom row). Posterior over the set of feasible manifold locations approximated using a Gaussian (red circle). At each step, random manifold locations are sampled from the posterior, and corresponding responses are reproduced using the artificial retina, and feedback from the subject is used to update the posterior using Monte Carlo methods (all samples in green, accepted samples in red). In ∼20 steps, the estimated manifold location (orange) converges to the true underlying location (blue).

These three approaches were examined using a model trained with 71 recordings and tested with 24 recordings. The proximity of manifold locations identified by approximation and optimization suggested that these approaches accurately capture properties of the new retina (Figure 4B). To examine the efficiency benefits of using the manifold, the optimization approach with limited data was examined in detail. The manifold location converged quickly as the recording duration increased, with ∼3 minutes of light response data producing a location similar to that produced by ∼30 minutes of data (Figure 4C). Although optimization of the manifold location predicted light responses with similar accuracy as training the full model (along with the CNN) when tested with a large amount of data (15-30 min), the manifold approach more accurately predicted light responses when the data were limited (∼8 sec) (Figure 4D).

The low dimensionality of the manifold enabled efficient generalization to a new retina, but in principle it could also reduce accuracy. On varying the manifold dimensionality, the ability to predict responses to previously unseen retinas saturated at ∼15 dimensions (Figure 4E, black line), a value that did not change with the number of retinas used for training (Figure 4E, colored lines). Hence, in addition to the previous observation on generalization to new stimuli within the collection of retinas used for learning (Figure 1D), the low-dimensional manifold is able to generalize in another way -- to new, previously unseen retinas.

These findings suggest that the manifold may aid in translating our understanding of the macaque retina to the human retina, an important goal for biomedical research. Recent work (Kling et al., 2020; Soto et al., 2020) has shown that the receptive field properties of the four numerically dominant RGC types (ON and OFF parasol and midget) are similar to those of their macaque counterparts. To test whether light responses in the human retina fall within the range observed across many macaque retinas, the manifold location of a single human retina was identified using the three operations described above (averaging, approximation, optimization). For each method, the estimated manifold location of the human retina was well within the span of manifold locations of many macaque retinas (Figure 4B,C), and the responses of a new retina were predicted with similar accuracy and efficiency in the two species (Figure 4D). Hence, the manifold reveals that the function of certain RGC types in the human retina can be understood as falling within the range of variation of macaque retinas.

Given that the macaque retinal code translates accurately to humans, it may provide a valuable tool for the development of an advanced artificial retina for treating vision loss. However, a challenging first step in restoring vision with such a device is to identify how the neurons in a blind retina should encode visual stimuli with the implant, based on the particular neural code of the individual receiving the implant. In this setting, the retina is no longer light-sensitive, so the healthy neural code cannot be identified directly. However, the human subject could potentially report the similarity of artificially induced images to a verbally described object. The neural encoding that produces the most accurate perception could then be identified by estimating the location of the retina on the low-dimensional manifold, using a psychophysical discrimination task. Such a task was simulated using the following iterative procedure: (i) sample a few of the plausible manifold locations, (ii) use each of these manifold locations to predict retinal responses for a particular visual stimulus, (iii) stimulate with the implanted device to produce each of these responses, (iv) ask the subject which stimulus produced a visual sensation that most closely matches a verbal description, and (iv) update the set of plausible manifold locations.

To illustrate the feasibility of this procedure, the above steps were simulated assuming (for simplicity) an artificial retina that has perfect cellular selectivity of stimulation. For updating the plausible locations, the perceptual accuracy was assumed to be governed by the Kullbeck-Leibler divergence between the distribution of neural responses associated with the tested manifold location and the distribution of responses associated with the true location. The set of plausible locations identified by the procedure converged to the true location (given by optimization; see above) in <20 small number of steps (see Methods, Figure 4F). Hence, the low-dimensional manifold could provide valuable efficiency in translational applications.

## Discussion

This work presented a new method to model individual variability in the neural code of the macaque retina using data from a large number of recordings. The approach employed a deep neural network that captured the shared response properties among retinas along with a low-dimensional manifold that captured differences between them. The manifold preserved the known invariances in neural encoding, while capturing the inter-dependence of response properties across cell types. It also revealed previously unknown sex-related differences in neural encoding, and provided more efficient ways of testing hypotheses on complex data sets in the face of both biological and experimental variability. Finally, the low-dimensional manifold enabled novel methods for identifying the neural code in a new recording, paving the way for exploiting large-scale datasets to optimize artificial vision with a retinal implant.

Note that, in the present data, biological and experimental variability for the most part could not be definitively separated. At least three sources of *in vivo* biological variability in the neural code could be present in the data: variation between animals, differences between the two eyes, and variation across retinal locations in a given eye. However, these sources of variability are inherently confounded with variation in three experimental procedures: euthanasia, eye removal, and *ex vivo* recording, respectively. In addition, some of the possible factors of inter-individual variation, such as the history of experiments and medical procedures throughout the lifespan of the animal, are neither entirely biological nor entirely experimental. Thus, although the relative importance of these three sources of variability could be studied further, for the most part, it is not possible to reliably isolate biological and experimental variation in the present data. An exception is the observed difference in retinal encoding between males and females: there were no known associations between the *ex vivo* experimental procedures used and the sex of the animal. Although there is no way to definitively exclude the possibility of experimental differences, these analyses strongly suggest that the male-female differences observed reflect true biological differences. Additionally, another biological factor, the proximity of dendritic lamination of specific cell types, is associated with *covariation* in measured response properties across preparations.

Importantly, the tools developed here make it possible to capture and analyze *both* experimental and biological variability in a single framework. This is an asset for understanding the neural code more completely using experimental data, which are always imperfect. For example, the low-dimensional manifold made it possible to project the data into a subspace orthogonal to known confounds, and thus to control for them without conditioning on specific experimental variables, thus retaining all the available data and statistical power.

The manifold framework can be used to understand biological and experimental variability in a principled manner. This is an extension of the idea that individual variability creates a manifold in the neural response space, with different manifolds corresponding to different possible invariances across recordings (Figure 2A). Isolated changes in a particular experimental or biological factor (while keeping the other factors fixed) could be conceptualized as producing a sub-manifold (Figure 5). Although the data used for this paper are not sufficient to study these sub-manifolds, we see evidence of additional structure within the manifold, such as the varying levels of covariation of response properties across cell-types and their relation to dendritic lamination.

**Figure 5:**
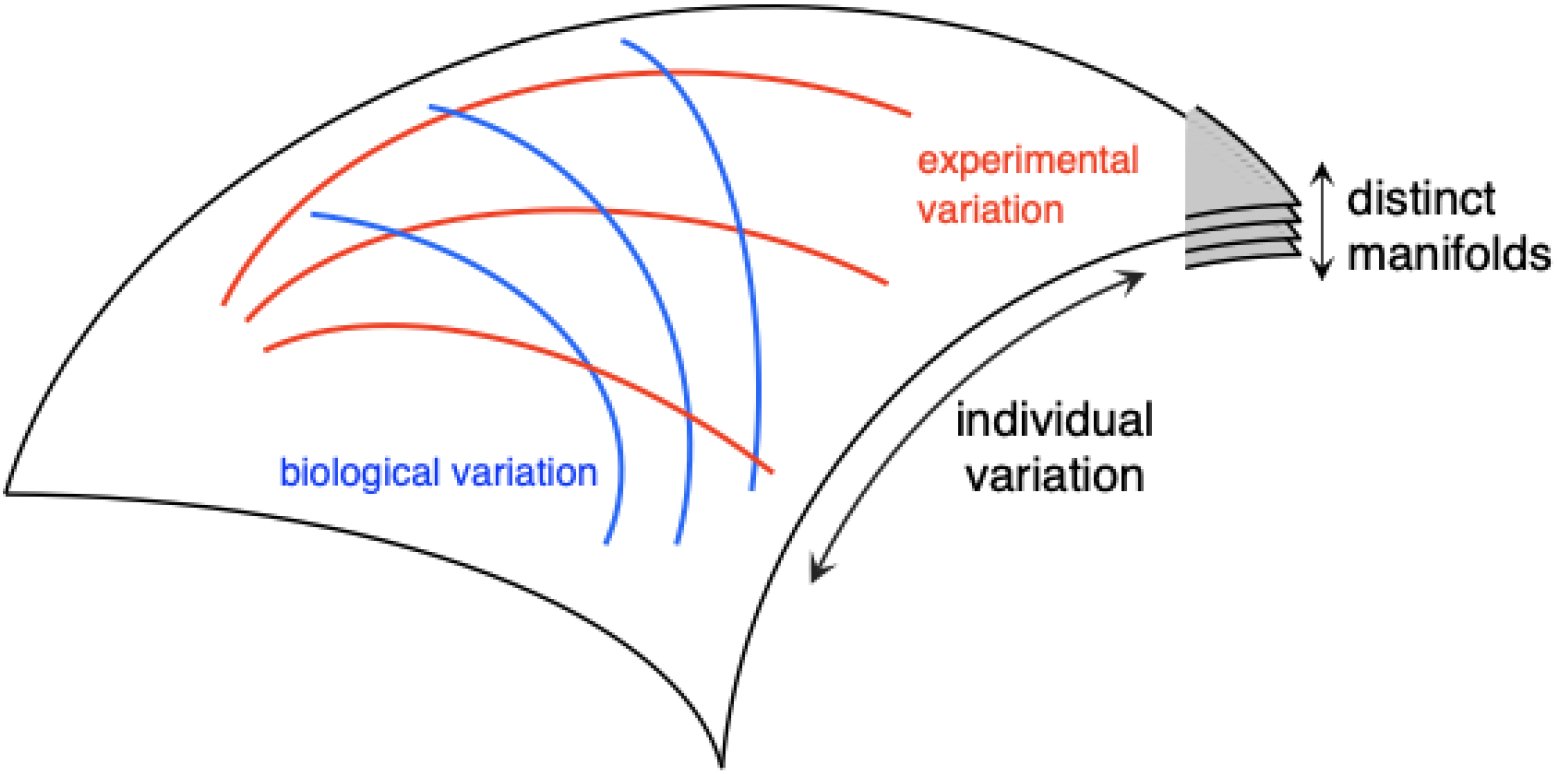
Sub-manifolds associated with variations in experimental and biological factors. Each submanifold is defined by the manifold locations spanned by the variation in one factor while keeping the other factors fixed. In the illustration, red (blue) curves correspond to variation in a particular experimental (biological) factor, while keeping other factors fixed. Different settings of the fixed factors correspond to a transformation of the submanifold (different curves) for a given varying factor.

For certain scientific questions it is not necessary to identify the underlying cause of variability. For example, attributing the varying degrees of covariation of different response properties (temporal kinetics, nonlinearity, autocorrelation) and cell types (ON vs. OFF; parasol vs. midget) to either biological or experimental factors is not crucial for understanding the dependence on a biological variable like lamination depth. For this observation, the presence of either biological or experimental variability can be understood as a “perturbation experiment” that reveals the interdependence of features of neural responses, which presumably has a biological origin.

The neural coding differences observed between male and female retinas add to a large literature on sex-based differences in brain structure and function (Cahill, 2006; Choleris et al., 2018; Poplin et al., 2018). In the retina, genetic differences between male and female primates (including humans) produce different variation in cone photopigment spectral sensitivities, and thus different color vision, across the population (National Eye Institute, 2019). However, to our knowledge, differences in neural coding between males and females have not been reported, perhaps due to the lack of appropriate physiological recordings and/or analysis tools.

Several recent studies have shown that neural responses in the retina, which have often been described using pseudo-linear models (Chichilnisky, 2001; Pillow et al., 2008), can be modeled more accurately using artificial neural networks (Batty et al., 2017; McIntosh et al., 2016). However, the function of these complex models can be difficult to interpret. In the present work, the machine learning model was used to more accurately capture shared (and complex) aspects of neural response, but information about variability between recordings was summarized in a simple, interpretable low-dimensional manifold (Schneidman et al., 2001). In principle, such an approach could be used for other applications in which a neural computation of interest is captured using a simple low-dimensional manifold, while a machine learning model improves accuracy by capturing the possibly complex aspects of the computation that are of less immediate interest.

The manifold of neural coding variability may also be useful in other neuroengineering applications, such as motor prostheses. The goal of a motor prosthesis is usually to read out the neural activity in a paralyzed person to control a computer cursor or a robotic limb (Bensmaia and Miller, 2014). Similar to the problem of identifying the neural encoding in a blind person, identifying the neural mapping in a paralyzed person is limited by the absence of simultaneous neural recordings and limb trajectory measurements, and thus may benefit from leveraging existing data that captures the diversity of neural coding across individuals. Specifically, a manifold of inter-individual variation may be useful for identifying neural decoding in a person with an implant, perhaps using a task involving imagined movements to identify the manifold location of a particular person (Shenoy and Carmena, 2014). The manifold may also be useful for dealing with the challenge of variability over time in chronic recordings (Chestek et al., 2011).

The neural coding manifold may also be useful for harnessing brain plasticity, which could improve vision with an artificial retina. Indeed, present-day retinal implants make little attempt to reproduce the neural code of the retina, and thus implicitly rely heavily on plasticity to compensate for device limitations (Beyeler et al., 2017). In motor prostheses, it has been shown that the brain can more easily adjust its activity to accommodate perturbations in the artificial neural decoder if these perturbations lie in a low-dimensional manifold (Golub et al., 2018). In the case of an artificial retina, we hypothesize that the brain may more readily learn to interpret the neural activity produced by the implanted device if the visual encoding that it uses is designed to lie within the manifold of retinal coding variability.

## Acknowledgements

We thank J. Carmena, K. Bankiewicz, T. Moore, W. Newsome, M. Taffe, T. Albright, E. Callaway, H. Fox, R. Krauzlis, S. Morairty, and the California National Primate Research Center for access to macaque retinas. We thank Kristy Berry, K. Williams, B. Morsey, J. Frohlich, and M. Kitano for accumulating and providing macaque retina metadata. The human eye was provided by Donor Network West (San Ramon, CA). We are thankful for the cooperation of Donor Network West and all of the organ and tissue donors and their families, for giving the gift of life and the gift of knowledge, by their generous donations. We thank K. Shenoy, G. Field, F. Rieke, W. Newsome, E. Simoncelli, S. Mitra, V. Gupta, K. Talwar, V. Feldman, A. Gogliettino, E. Wu, S. Madugula, M. Zaidi and the entire Stanford Artificial Retina team for helpful discussions. We thank Google internship and Student Researcher programs (NPS), Research to Prevent Blindness Stein Innovation Award, Wu Tsai Neurosciences Institute Big Ideas, NIH NEI R01-EY021271, NIH NEI R01-EY029247, and NIH NEI P30-EY019005 (EJC), NSF IGERT Grant 0801700 (NB) and NIH NEI F31EY027166 (CR), NSF GRFP DGE-114747 (NB and CR) for funding this work.

## Methods

### Recordings

Preparation and recording methods are described elsewhere (Chichilnisky and Kalmar, 2002; Frechette et al., 2005; Litke et al., 2004). Briefly, eyes were enucleated from terminally anesthetized macaque monkeys (*M. Mulatta* or *M. Fascicularis*) used by other experimenters in accordance with institutional guidelines for the care and use of animals. Immediately after enucleation, the anterior portion of the eye and the vitreous were removed in room light. A human eye was obtained from a brain dead donor (29 year-old Hispanic male) through Donor Network West (San Ramon, CA) (See (Kling et al., 2020) for details.)The eye was stored in darkness in oxygenated Ames’ solution (Sigma, St. Louis, MO) at 33°C pH 7.4. Segments of isolated or RPE-attached peripheral retina (approximately 3mm x 3mm, taken from 6-15mm temporal equivalent eccentricity(Chichilnisky and Kalmar, 2002)) were placed flat, RGC side down, on a planar array of 512 extracellular microelectrodes arranged in an isosceles triangular lattice. The electrode spacing was 60μm in each row, and the array covered a rectangular region measuring 1800 µm x 900 µm. While recording, the retina was perfused with Ames’ solution (31-36°C; typically 32 °C for RPE attached and 34°C for RPE isolated dissections), bubbled with 95% O_2_ and 5% CO_2_, pH 7.4. Voltage signals on each electrode were bandpass filtered (80Hz - 2kHz), amplified, and digitized at 20 kHz (Litke et al., 2004).

A custom spike-sorting algorithm was used to identify and segregate spikes from distinct cells(Litke et al., 2004). Briefly, candidate spike events were detected using a threshold on each electrode, and voltage waveforms on the electrode and nearby electrodes in the 4ms period surrounding the time of the spike were extracted. Candidate neurons were identified by clustering the waveforms using a Gaussian mixture model. Candidate neurons were retained only if the assigned spikes exhibited a 1 ms refractory period and had a stable firing rate for the entire duration of recording. Duplicate spike trains were identified by temporal cross-correlation and removed. For each cell, the autocorrelation function of the recorded spike train was computed by computing the correlation coefficient for different time lags.

### Visual stimuli and cell type identification

Visual stimuli were delivered using the optically reduced image of a CRT monitor refreshing at 120 Hz and focused on the photoreceptor outer segments. The optical path passed through the transparent plug and Ames’ solution or through the mostly transparent electrode array and the retina. The relative emission spectrum of each display primary was measured with a spectroradiometer (PR-701, PhotoResearch) after passing through the optical elements between the display and the retina. The total power of each display primary was measured with a calibrated photodiode (UDT Instruments). The mean photoisomerization rates for the cone photoreceptors were estimated by computing the inner product of the primary power spectra with the spectral sensitivity of each cone type, and multiplying by the effective collecting area of primate cones (∼0.6 µm^2^)(Angueyra and Rieke, 2013; Schnapf et al., 1990), resulting in photoisomerization rates of approximately 800–2200, 800–2200, 400–900 for the long-, middle- and short-wavelength sensitive cones, respectively. The stimulus pixel size on the retina was either 41.6 microns (8 monitor pixels), 52 microns (10 monitor pixels) or 83.2 microns (16 monitor pixels). A new white noise frame was drawn at refresh rates of 60 Hz or 30 Hz. The pixel contrast (difference between the maximum and minimum intensities divided by the sum) was 96% for each display primary, with mean intensity of 50%. The white noise stimulus either modulated the three display primaries independently, or coherently, at each spatial location.

In each recording, RGCs were classified into distinct types using properties of the spatial receptive field and response time course obtained from the spike-triggered average (STA) stimulus (Chichilnisky, 2001; Chichilnisky and Kalmar, 2002; Field and Chichilnisky, 2007). A two-dimensional Gaussian fit to the spatial receptive field was used for determining the center location(Chichilnisky and Kalmar, 2002). All analyses used recordings with stable firing rates and nearly complete tiling of ON and OFF parasol cell receptive field mosaics. For Figure 3, only recordings that also had nearly complete ON and OFF midget cell mosaics were used. For model fitting, both the visual stimulus and spike times were binned at 8.33ms (120Hz), and the visual stimulus was upsampled to 8 monitor pixels, resulting in a common 80×40 pixel grid across recordings.

### Linear Nonlinear Poisson model

The Linear Nonlinear Poisson (LNP) model consists of a linear spatio-temporal filter followed by a point nonlinearity(Chichilnisky, 2001). A filter that is separable in space and time was used(Chichilnisky and Kalmar, 2002), which is equivalent to a cascade of a spatial filter and a temporal filter. These filters were estimated for each cell in each recording as follows. First, the STA was computed by averaging the stimulus preceding spikes, over all pixel locations and 250ms (30 frames at 120Hz) prior to the spike. Next, the spatial filter was computed by choosing the STA frame with the single largest pixel magnitude. The spatial filter was restricted to a rectangular window around the receptive field. The receptive field was defined as the set of pixels with absolute magnitude greater than 2.5σ, contiguous with the strongest pixel, where σ is the robust standard deviation(Rousseeuw and Croux, 1993) of pixels in the STA, an estimate of the measurement noise. Next, the temporal filter was identified by averaging the time course of all pixels in the receptive field. Finally, the output nonlinearity was estimated by fitting a 5th order polynomial to the relationship between the observed responses and the generator signal, which was computed by filtering the stimulus with the estimated spatial and temporal filters(Chichilnisky, 2001).

### Neural network model

A convolutional neural network was used to predict RGC responses across multiple recordings simultaneously. Below, the model architecture and the fitting procedure are described in detail.

#### Notation

The model *f*(*S*, α_*i*_, *C*_*i*_), takes as its input the visual stimulus *S*, recording-specific information about the collection of recorded cells *C*_*i*_, and the recording-specific manifold location α_*i*_, and yields as its output the predictions for recording-specific response *R*_*i*_. For simplicity, the following description is for a single sample stimulus-response pair, but it can be extended to multiple stimuli and corresponding responses in a straightforward manner.

The recent history of the visual stimulus is given by 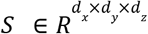, where *R* is the set of real numbers, *d*_*x*_ × *d*_*y*_ are the spatial dimensions (80 x 40), and *d* is the number of time bins (30). Stimuli presented at different spatial or temporal resolutions were upsampled or downsampled to these dimensions.

The recording specific manifold location is given by α ∈ *R*^*n*^, where *n*is the manifold dimensionality.

The recording specific information about recorded cells is given by 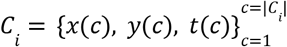, where each cell *c* is described by its receptive location (*x*(*c*), *y*(*c*)) in the *d*_*x*_ × *d*_*y*_ visual space and its cell type *t*(*c*). For models with only two cell types, *t*(*c*) ∈ {0, 1}, for ON and OFF parasols respectively. For models with four cell types, *t*(*c*) ∈ {0, 1, 2, 3}, corresponding to ON parasol, OFF parasol, ON midget, and OFF midget cell types.

The responses are given by 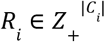, where *Z*_+_ denotes non-negative integers and |*C*_*i*_ | is the number of cells in recording *i*. Responses were binned at the same resolution as the stimulus (8.33ms).

#### Model architecture

The model *f*(*S*, α_*i*_, *C*_*i*_) passes the visual stimulus *S* through a multilayered convolutional neural network, with each layer consisting of a convolution (stride 1), retina-specific normalization and a point-wise (softplus) nonlinearity (see Figure 1B). The model output is Poisson firing rate. This firing rate is used to predict the responses *R*_*i*_. The number of channels and filter sizes are chosen by cross-validation, as described below. Recording-specific normalization and challenges associated with predicting responses for varying numbers of cells across recordings are also given below.

Recording-specific normalization is inspired by previous work(Dumoulin et al., 2017), in which a translationally-invariant affine transformation of the layer activations adapts the model to each recording. The scale and shift coefficients for this affine transform are determined linearly from the manifold location α_*i*_. Let 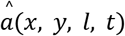 be the activation after convolution at location *x, y* in the channel *l* of layer *t*. First, the mean μ and standard deviation σ across samples in a batch are computed, and used to calculate normalized activations: 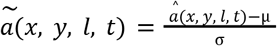. Next, using the manifold location α_*i*_, a learned affine transform determines the desired mean 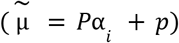 and standard deviation 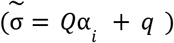 for each layer. Finally, the normalized activations are transformed to give recording-specific activations 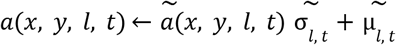. Note that the retina-specific scales and shifts are the same for each location in visual space, preserving the translational invariance of convolutional networks and reflecting the homogenous response properties of the RGCs belonging to a single type.

A potential challenge is that the number of recorded neurons, and hence the number of outputs of *f*(.), is variable across recordings. To address this issue, the model predicts multiple response maps, one for each cell-type, with the same spatial dimensions as the visual stimulus. The response for each cell is read off from its cell location in the response map of the corresponding cell type. Specifically, *f*(.)outputs *m*_*i*_ (*x, y*), which corresponds to firing rate map of cell-type *i*, and for a cell with type *t*(*c*)and centered at *x*(*c*), *y*(*c*), the Poisson firing rate is given by *m*_*t*(*c*)_ (*x*(*c*), *y*(*c*)).

#### Model fitting

Estimation of recording-specific parameters (α_*i*_)and the shared parameters are performed by maximizing the log-likelihood of observed responses, summed across all the cells, recordings and stimuli. This is performed by stochastic gradient descent, where at each step, a randomly sampled batch of stimuli and corresponding responses from a particular recording are used to update the shared and the corresponding recording-specific parameters. The batch size was 250 and the updates were performed using the Adam(Kingma and Ba, 2014) update algorithm with learning rate of 0.1. For each training retina, the first 4 min of white noise data were used for testing and the remainder was used for training. The duration of the stimulus varied from 15-90 min (median 30 min) across experiments. A model with 4 layers, 3 x 3 or 1 x 1 filter size, 64 channels per layer and a 15 dimensional manifold was chosen based on cross validation and used for subsequent analysis (see Figure 1B for architecture).

### Variation of neural coding on the manifold

The following steps were used to test if the manifold captured variations in neural response properties across recordings. The estimated manifold locations were similar when estimated from each smaller partition at the beginning and end of the entire recording, suggesting a stable manifold for further analysis. First, the manifold direction that was maximally correlated to the variations of a particular response feature was identified by linear regression. Second, recordings were projected along this direction, and the Spearman rank correlation with the response property was measured. Statistical significance was measured with a permutation test, where the null distribution was generated by permuting the recordings with the manifold locations fixed. In Figure 2, the Spearman rank correlation and its statistical significance was measured for the first principal component projection of various response properties such as receptive field size (ON: 0.78, p<0.0001 ; OFF: 0.76, p<0.0001), time course (ON: 0.83, OFF: 0.85; p < 0.0001), output non-linearity of the LNP model (Spearman rank correlation for ON: 0.92; OFF: 0.92; p < 0.0001) and auto-correlation (ON: 0.84, p<0.001 OFF: 0.62, p < 0.05). The interdependence between response properties was either measured directly in raw data using Spearman rank correlation, or with the angle between the corresponding manifold directions. Pairwise correlations between all ten response properties considered (firing rate, output nonlinearity, time course, autocorrelation & RF area across ON and OFF parasols) was captured in a ∼6 dimensional space (90% of variance captured).

### Relating the manifold location to biological and experimental factors of variation

The estimated model was examined to see the dependence of manifold location on various biological and experimental factors of variation. For analysis of sex differences, only the subset of recordings (102) from one species (M. Mulatta) were used. Based on the documented medical history, the animals with a gender specific condition were not considered for this analysis. First, the separation between male and female recordings was measured by computing the d’ value of the projection of the two distributions onto the difference in the means. The d’ value observed (∼1.8) indicated that the sex based differences were not large on an individual basis. Second, a bootstrap test was performed to test whether the mean locations of the male and female recordings were statistically distinguishable. The distance between the mean manifold locations of male and female retinas was measured and compared to a null distribution of distances generated by resampling (with replacement) of the manifold locations. The null distribution was fitted with a normal distribution and the significance level was measured as the probability mass greater than the observed distance in data. Because multiple recordings were frequently recorded from the same animal, a hierarchical variant of this bootstrap test was performed, in which resampling was performed according to the hierarchical structure of the data (Saravanan et al., 2019), by first sampling an animal and then sampling the manifold location of one of the recordings from that animal, both with replacement. Hierarchical bootstrap is more conservative and biased towards accepting the null hypothesis (Saravanan et al., 2019). Mean manifold locations for male and female retinas were significantly different (p<10^−6^ for bootstrap and p<0.05 for hierarchical bootstrap). Identical tests were applied for assessing male-female differences in firing rate and the speed of temporal filtering (p<0.01 for bootstrap and p<0.05 for hierarchical bootstrap for both quantities).

Apart from male-female differences, systematic differences in visual encoding properties were not observed with respect to animal age, laterality of the eye, temporal/nasal location, circadian time of dissection, month/season of the experiment and the primate lab from where the retina was obtained. Note that it is important to be careful when testing multiple statistical hypotheses using the same dataset. For the present data, the male-female differences were highly significant (p<10^-6) using the marginalization method, even after comparing with a lower Bonferroni-corrected threshold corresponding to multiple (∼10) hypothesis tests.

### Invariance of neural coding on the manifold

The ability of the manifold to preserve previously reported invariances of the neural code was tested as follows. First, random manifold locations were sampled by perturbing the learned locations of training retinas with a Gaussian noise of standard deviation equal to their median nearest-neighbor distance. Second, a ∼800 sec long white noise stimulus was sampled, and ON and OFF parasol firing rate maps were computed using the neural network, which was adapted using the manifold locations. Third, the Poisson firing rates for 200 cells (100 of each type) with random receptive field locations were read off from the firing rate maps. Finally, the cell responses were sampled, and used to estimate a Linear-Nonlinear Poisson model, which served as an interpretable summary of neural encoding captured by the manifold location. Comparison of average receptive field size and the zero crossing time of the temporal filter revealed known invariances between ON and OFF parasols (Figure 2J, K).

### Estimation of the manifold location of a previously unseen retina

By fixing the shared parameters after learning, and estimating the recording-specific representation on the manifold, the trained model was adapted to predict responses in a new, previously unseen recording. Based on the amount of data available, several methods can be employed to identify the manifold location (Figure 4). These methods are described below in detail.

#### Averaging

When no data about the new retina are available, the simplest approach is to average the locations of all the retinas used for training.

#### Approximation

This is similar to averaging, but only using the subset of training retinas with similar response properties as the new retina. For Figure 4, locations of five training retinas with the most similar firing rate function were used for approximation.

#### Optimization

When light response data are available, the manifold location α_*i*_ of retina *i* was determined by Bayesian inference. Bayesian inference combines a Gaussian prior (*P*_*prior*_ (α) ∼ *N*(μ_*prior*_, σ _*prior*_)) over manifold locations determined from the training retinas and the likelihood 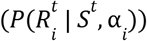 of collection of stimulus-response data for the new retina. The posterior 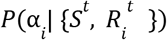 was maximized using gradient ascent (learning rate 0.1) :

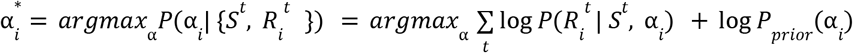

#### Discrimination task

When light responses are unavailable, as in the case of a blind person implanted with a retinal prosthesis, manifold location could instead be estimted using a discrimination task. The effectiveness of this proposed approach was validated in simulation. For a given visual stimulus, the discrimination task involves using the implanted retinal prosthesis to reproduce responses corresponding to two manifold locations and asking the subject to select the response that yields perception most closely matching a verbally described stimulus. Multiple rounds of this task are used to update the posterior on manifold locations.

The discrimination task was simulated under the assumption that the perceptual difference of the responses generated by hypothetical retinas at two manifold locations α_1_ and α_2_ for a stimulus *S* is equal to the KL-divergence between the corresponding response distributions *P*(*R*|*S*, α_1_) and *P*(*R*|*S*, α_2_). Given α_*true*_ as the true underlying manifold location, the blind person’s feedback*Y*(α_*true*_, α_1_, α_2_) = 0 if

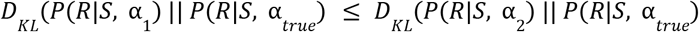

and *Y* = 1 otherwise. For simplicity, sampled responses were used to compute an unbiased estimate of the KL-divergence :

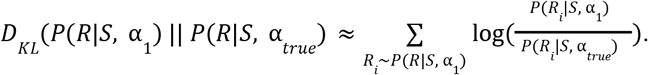

Hence, the posterior over manifold location after *t* steps of the task is given by:

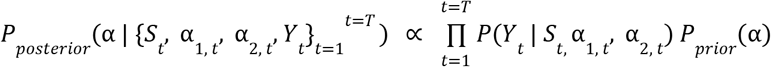

where the prior is estimated from training retinas as a Gaussian distribution: *P*_*prior*_ (α) ∼ *N*(μ_*prior*_, σ_*prior*_). In the simulations, the visual stimulus *S*consisted of letters of English alphabet, flashed for 100 *ms*and preceded and succeeded by 50 *ms*of gray screen. At each step, a Gaussian approximation of theposterior *P*_*posterior*_ (α) ∼ *N*(μ_*posterior*_, σ_*posterior*_) was maintained, and updated using Monte-Carlo sampling. In summary, the steps for the ^*th*^iteration of the algorithm are as follows :

1. Sample symmetric α_1,*t*_, α_2,*t*_ around posterior mean: α_1,*t*_ ∼ *P*_*posterior*_ (α); α_2,*t*_ = 2μ_*posterior*_ − α_1,*t*_.
2. Sample an English letter and a target stimulus *S*_*t*_.
3. Sample responses *R*_1,*t*_ ∼ *P*(*R*|*S*_*t*_, α_1,*t*_); *R*_2,*t*_ ∼ *P*(*R*|*S*_*t*_, α_2,*t*_).
4. Get patient feedback *Y*(α_*true*_, α_1,*t*_, α _2,*t*_), based on an estimate of the KL divergence using sampled responses *R*_1,*t*_, *R*_2,*t*_.
5. Update the posterior of plausible manifold locations.
  a. Sample *N* retina locations α_*i*_ ∼ *P*_*posterior*_ (α) for *i* ∈ [1, …, *N*].
  b. For the set of sampled manifold locations, find the subset that matches user feedback, i.e., with *Y*(α_*i*_, α_1,*l*_, α_2,*l*_) =*Y*(α_*true*_, α_1,*l*_, α _2,*l*_)for all *l* (= 1,…, *t*)previous steps. Let this subset of be 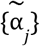.
  c. Update posterior distribution with 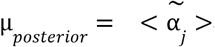 and 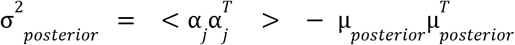.

For results shown in Figure 4F, α_*true*_ was set as the result of optimizing the manifold location using light-evoked responses. In the simulations, the posterior distribution converged in ∼20 steps, suggesting that the low dimensional manifold can be used for efficiently identifying the expected neural code in a blind person. However, the amount of noise in the simulation is probably lower compared to what would be encountered in practice, leading to a larger number of steps to identify the true manifold location and may perhaps require changes to the estimator of KL divergence and the method to update the posterior of α.

## Data/Code Availability

The data/code that support the findings of this study are available from the corresponding author upon reasonable request.

